# From motor protein to toxin: Mutations in the zonula occludens toxin (Zot) of *Vibrio cholerae* phage CTXϕ suggest a loss of phage assembly function

**DOI:** 10.1101/2022.09.21.508815

**Authors:** Long Ma, Simon Roux, Xiaoting Hua, Yong Wang, Belinda Loh, Sebastian Leptihn

## Abstract

Prophages, i.e. dormant viruses residing in bacterial cells, are not just passive passengers in the bacterial host. Several prophage-encoded genes have been shown to be contributors to bacterial virulence by mediating antimicrobial resistance or by providing toxins. Other prophage genes exhibit beneficial effects on the host by modulating e.g. motility or biofilm formation. In this study, we used an *in vivo* phage assembly assay and tested an extensive array of single point mutations or their combinations found in Zot, the zonula occludens toxin encoded by the *Vibrio cholerae* phage CTXϕ. The assay makes use of the highly homologous Zot-like protein g1p of the filamentous Coliphage M13, a motor protein that mediates the trans-envelope assembly and secretion of filamentous phages. We also measured the *in vitro* ATP hydrolysis of purified proteins, and quantified virus production in *V. cholerae* mediated by Zot or the Zot-like protein of the two *Vibrio* phages CTXϕ and VFJϕ. In addition, we investigated sequence variations of the Walker motifs in *Vibrio* species using bioinformatics method, and revealed the molecular basis of ATP binding using molecular docking and molecular dynamics simulation based on the structure predicted by AlphaFold2. Our data indicates that g1p proteins in *Vibrio* can easily accumulate deleterious mutations and likely lose the ability to efficiently hydrolyse ATP, while the CTXϕ Zot was further exapted to now act as an auxiliary toxin during the infection by *Vibrio cholerae*.

## Introduction

Prophages, which exist in a “dormant” state until activation and initiation of replication, can have a substantial impact on the host. A small minority of prophage-encoded genes are not directly required for the viral replication cycle or genome insertion, nor do they encode for structural components of the phage particle. Some of these “prophage accessory” genes have been identified as virulence factors such as antimicrobial resistance genes or toxins. Prominent examples are the Shiga toxin- and the Cholera toxin-encoding genes ^1–4^. Other genes appear to influence bacterial host virulence indirectly by e.g. inducing motility or biofilm formation. Similar to other prophages, filamentous phages, which lead to a “chronic infection” with continuous viral replication while not killing the host, can play an important role in the pathogenicity and fitness of their hosts ^5,6^. While some inoviruses have been observed to have a negative impact on fitness of infected bacteria, virulence factors that are transmitted via infection by the virus or passed on vertically as a prophage, help the host to obtain an evolutionary advantage ^7^. One of the best examples is the CTXφ phage infecting *Vibrio cholerae* that has been shown to be responsible for transmitting the genes coding for the cholera toxin CTX to its host, rendering it highly pathogenic in widespread cholera epidemics ^8^.

Filamentous phages are Inoviruses, single-stranded DNA viruses, with an extraordinary morphology being essentially protein coated DNA filaments at approximately 1 μm in length and 6 nm in diameter ^9^. In contrast to the head-and-tail phages, filamentous phages are less common and are often thought of as an oddity among the prokaryotic viruses. However, they are found in almost all bacteria and possibly also archaea ^10^, and particularly those in pathogens are of interest with regards to their contribution to virulence of their hosts ^6,11–13^. Filamentous phages are non-lytic and are extruded from the bacterial envelope as the host continues to grow. After infecting their host, filamentous phages remain as an episomal element or integrated into the genome while the phage DNA is being replicated and viral particles are being produced, a “chronic infection”. The genomes of *Vibrio cholerae* contain a plethora of prophages, including those of filamentous phages. Interestingly, according to a previous report, so far not a single *V. cholerae* strain has been isolated that contains only a single chromosomally integrated filamentous phage ^14^. For genome integration, a single dif site in *Vibrio cholerae* is used by several phages including CTXφ and satellite phages RS1φ and TLCφ ^15,16^. The satellite phages RS1φ and TLCφ may require helper phages, as they do not encode morphogenesis proteins. In the case of RS1φ, the phages CTXφ or KSF-1φ were proposed to mediate the encapsidation of the RS1φ genome, similar to TLCφ mediating the formation of fs2φ viral particles ^17–20^.

The archetypical filamentous phages fd and M13 infect *E. coli*, with a genome that encodes only 11 genes, devoid of any virulence genes and only contain those responsible for replication or morphogenesis. The products of five of these genes encode structural proteins that eventually cover the DNA strand in the mature virion. The remaining six genes encode proteins that are involved in the reproduction of genomic DNA in the host during the virus life cycle. Gene 1 is a phage-encoded gene whose protein is essential for the assembly and extrusion of the phage, and is referred to as g1p, gp1 or p1. The protein is also called a Zot-like protein, as it is homologous to the *zonula occludens toxin* of the filamentous *Vibrio cholerae* phage CTXϕ. One of the key features of gene 1 (g1p) is that it contains an internal ORF (gene 11, g11p) that has been shown to be necessary for phage production, and a transmembrane domain that anchors the complex to the inner membrane of the host ^21^. An essential feature of gene 1 is the presence of Walker motifs which are the molecular basis for ATPase activity ^22^.

Walker motifs are considered fundamental motifs in biological systems. Proteins found to possess consensus Walker A motifs, A/GxxxxGKT/S (where x represents any amino acid) and consensus Walker B motifs, hhhhDE (where h represents any hydrophobic residue) are highly likely to be functional ATPases ^23^. Previous work conducted in our lab has shown that the putative Walker A and Walker B motifs located at the N-terminus of g1p are required for the production of M13 ^22^. Sequence alignments of gene 1 have shown that the Walker motifs are present across most species of filamentous phages. A striking exception is found in the filamentous phage CTXϕ, which infects *Vibrio cholera* and has been shown to be responsible for the bacteria’s pathogenicity, as it supplies the host with the phage-encoded cholera toxin (CTX). The Walker motifs found in gene 1 of the CTXϕ phage have mutations in key positions in the consensus sequence that would theoretically render the protein inactive for ATPase activity. The CTXϕ protein Zot was investigated as one of two auxiliary toxins, in addition to CTX. Here, the Zot protein was tested on human cells and was found to increase the permeability of small intestinal mucosa by opening intercellular tight junctions ^24,25^.

In this work, we analysed the Zot/ Zot-like assembly genes of two filamentous *V. cholerae* phages and compared it with the model phage M13. While the Walker motifs in *V. cholerae* phage VFJϕ has a sequence in line with the consensus motif, CTXϕ exhibits a deviant Walker A motif and an uncommon Walker B motif, with bulky phenylalanine residues. Using a combination of protein biochemistry, bioinformatics and *in vivo* methods, we could demonstrate that the Zot of CTXϕ is highly inefficient in ATP hydrolysis and phage assembly.

## Materials & Methods

### Molecular biology

Seamless cloning was used to generate plasmids; Supplemental Table 1 lists mutants and chimeras of gene 1 (UniProtKB: P03656) and ZOT/ZOT-like genes VFJXϕ Zot (UniProtKB: R9TFZ4) and CTXϕ Zot (UniProtKB: P38442). QuikChange II site-directed mutagenesis was performed according to the company’s instructions (Agilent Technologies Inc., Santa Clara, CA, USA). The numbering of amino acids follows the sequence of the M13 g1p.

### *In vivo* complementation assays (Spot tests)

Testing point mutations *in trans* was conducted as previously described ^22^. Briefly, LB agar plates were top-layered with exponentially growing *E.coli* expressing protein, mixed with LB agar (0.7% agar). Once the agar solidified, dilutions of the M13 phage was then “spotted” and the plates were incubated at 37°C overnight to develop plaques. To quantify phage titer, dilutions of the supernatant were mixed with *E. coli* mixed with LB agar (0.7% agar). After incubating at 37°C overnight, plaques grown were counted, and the phage titer was calculated based on the dilution factor.

### Quantification of viruses in bacterial culture supernatant

To determine the exact amount of virus particles assembled in the case of *V. cholerae* phage, we performed qPCR experiments as “spotting” (described above) did not result in the formation of visible plaques-like zones. We designed three primer pairs based on the receptor-binding proteins of the phages (called g3p, gp3 or p3) (Supplemental Table 2) allowing to correlate the amount of M13 phages detected in our *in vivo* assay, using spotting, with the qPCR data. qPCR data was also correlated with the number of bacterial cells (to obtain phage production values per host cell), determining the number of bacteria by qPCR using the 16S ribosomal RNA gene (rrsA) ^26^. The *V. cholerae* strain we used had the CTXb gene and ZOT genetically removed from the CTXϕ (kind gift from Prof. Menghua Yang, Zhejiang A & F University, Hangzhou, China). CTXϕ ZOT and the ZOT-like genes from VFJϕ were cloned into pBAD33 and chloramphenicol-resistant colonies were selected after introduction of the respective plasmids by electroporation. Liquid cultures were grown until an OD600 of 0.2 was reached, then 0.5 μg/ml mitomycin C (final concentration) for prophage induction and 0.2% L-Arabinose (final concentration) for plasmid-derived protein expression, were added. After culturing for 4 hours, sedimentation and repeated washing, bacterial DNA was extracted from 1 mL using the BMamp Rapid Bacteria DNA kit (Biomed, catalog number: DL111-01) following the instruction manual. Next, we determined the number of phages in the supernatant. To 200 μL of supernatant 1 μL DNase (New England Biolabs, catalog number: M0303S) 1 μL RNaseA (Biomed, catalog number: 756780AH) was added and incubated at 37 °C for 5 hours, followed by heat denaturation at 70° C for 10 minutes to deactivate the enzymes. To remove proteins, 25 μL proteinaseK was added followed by DNA extraction using the PureLink Viral RNA/DNA Mini Kit (Thermo Fisher Scientific catalog number:12280050) according to the manufacturer’s instructions. For the qPCR we used the BlasTaq 2X qPCR MasterMix kit (Abm, catalog number: G891) in triplicates, with four independent experiments.

### ATP Hydrolysis assay

Expression and purification attempt of the entire genes were unsuccessful either due to toxicity to the cells, expression level or inability to capture the protein on the column, depending on the gene. Thus, only the ATPase domain containing cytoplasmic fragment of CTXϕ ZOT and the ZOT-like genes from VFJϕ were cloned into the *E. coli* expression vector pQE60 and expressed at 20°C over night to obtain C-terminal MBP-fusion proteins in M15 cells. Protein purification was performed with an AKTA pure system using MBPTrap HP 1ml column (Cytiva, catalog number: 28918778), protein concentration was determined via Bradford assay (Beyotime, catalog number: P0006). EnzChek Phosphate Assay Kit (Thermo Fisher Scientific, catalog number: E6646) was used to determine ATP hydrolysis rates according to the instructions provided by the manufacturer. After establishing a phosphate standard curve, the assay was performed with different amounts of purified proteins in triplicates with three independent repeats each using a newly expressed and purified protein batch.

### *In silico* sequence analyses

Phylogenetic Tree and sequence alignments were conducted using Clustal Omega with the output using ClustalW alignment format ^27^. Phylogenetic tree was visualised using iTOL ^28^.

### Structure prediction, molecular docking and molecular dynamics simulation

AlphaFold (version 2.2) with default pipeline was used to predict the structures of M13-g1p, VFJϕ-Zot, and CTXϕ-Zot. Five structures were predicted for each protein and the top one was subsequently docked with an ATP molecule in the cytoplasmic domain using AutoDock Vina ^29^. For each complex, six docked models were prepared.

The top ranked structure of the ATP-bound M13-g1p complex was embedded into a flat POPC lipid bilayer and solvated in a cubic water box containing 0.15 M NaCl, using the “Membrane Builder” function of the CHARMM-GUI webserver ^30^. The OPM (Orientations of Proteins in Membranes) webserver was used to align the transmembrane region (residue 254-270) in the lipid bilayer ^31^. The size of the simulated box was 12.0 nm, 12.0 nm and 13.9 nm in the x, y and z dimension, respectively, resulting in ~188,000 atoms in total. The CHARMM36m force field was used for the protein and the CHARMM36 lipid force field was used for the POPC molecules ^32^. ATP molecule was assigned with CHARMM CGenFF force field. TIP3P model was used for the water molecules. The system was then energy minimized and equilibrated in a stepwise manner using 1 ns NVT simulations and a following NPT simulation. Finally, a 200 ns productive simulation was performed. Neighbor searching was performed every 20 steps. Neighbor searching was performed every 20 steps. The PME algorithm was used for electrostatic interactions with a cut-off of 1.2 nm. A reciprocal grid of 100 × 100 × 112 cells was used with 4th order B-spline interpolation. A single cut-off of 1.2 nm was used for Van der Waals interactions. The temperature was kept at 310 K with the V-rescale algorithms. The pressure was kept at 1.0 × 10^5^ Pa with the Parrinello-Rahman algorithm. All simulations were performed using a GPU-accelerated version of Gromacs 2021.5 ^33^. Protein structures were visualized with PyMOL ^34^.

### Identification of Walker sequences in *Vibrio* genomes

1,529 *Vibrio cholerae* genomes were downloaded from NCBI RefSeq & GenBank databases (i.e. including potential redundancy) on June 28 2021 ^35^. Hmmsearch v ^36^ was used to compare all proteins predicted in these genomes to the g1p-like HMM profiles previously built from an extended catalog of inoviruses ^37^. *Vibrio cholera* g1p-like proteins were identified based on hits to these HMM profiles with a score ≥ 50.

## Results

### CTXϕ Zot phylogenetic separation from homologs of other phages indicate a divergent evolutionary pathway

While filamentous phages have mainly been found to infect Gram-negative bacteria, many have been identified in the genomes of most bacterial families and archaea ^37^. When analysing the amino acid sequence of Zot from several characterised filamentous phages, the sequence clearly aligns with other Zot or Zot-like proteins. However, several distinct features such as extended non-aligning stretches are found in the Zot gene of phage CTXϕ. The assembly proteins of M13 clusters with other proteins from Enterobacteriaceae (e.g. *Salmonella enterica, Klebsiella pneumoniae*) but also with *Vibrio cholerae* phage VFJϕ, while CTXϕ Zot exhibits a larger evolutionary distance to the cluster (Figure 1A). This might indicate that the sequence has undergone evolutionary changes and may have evolved to possibly adopt a structure (and function) different from that of a phage assembly motor.

**Figure 1:**
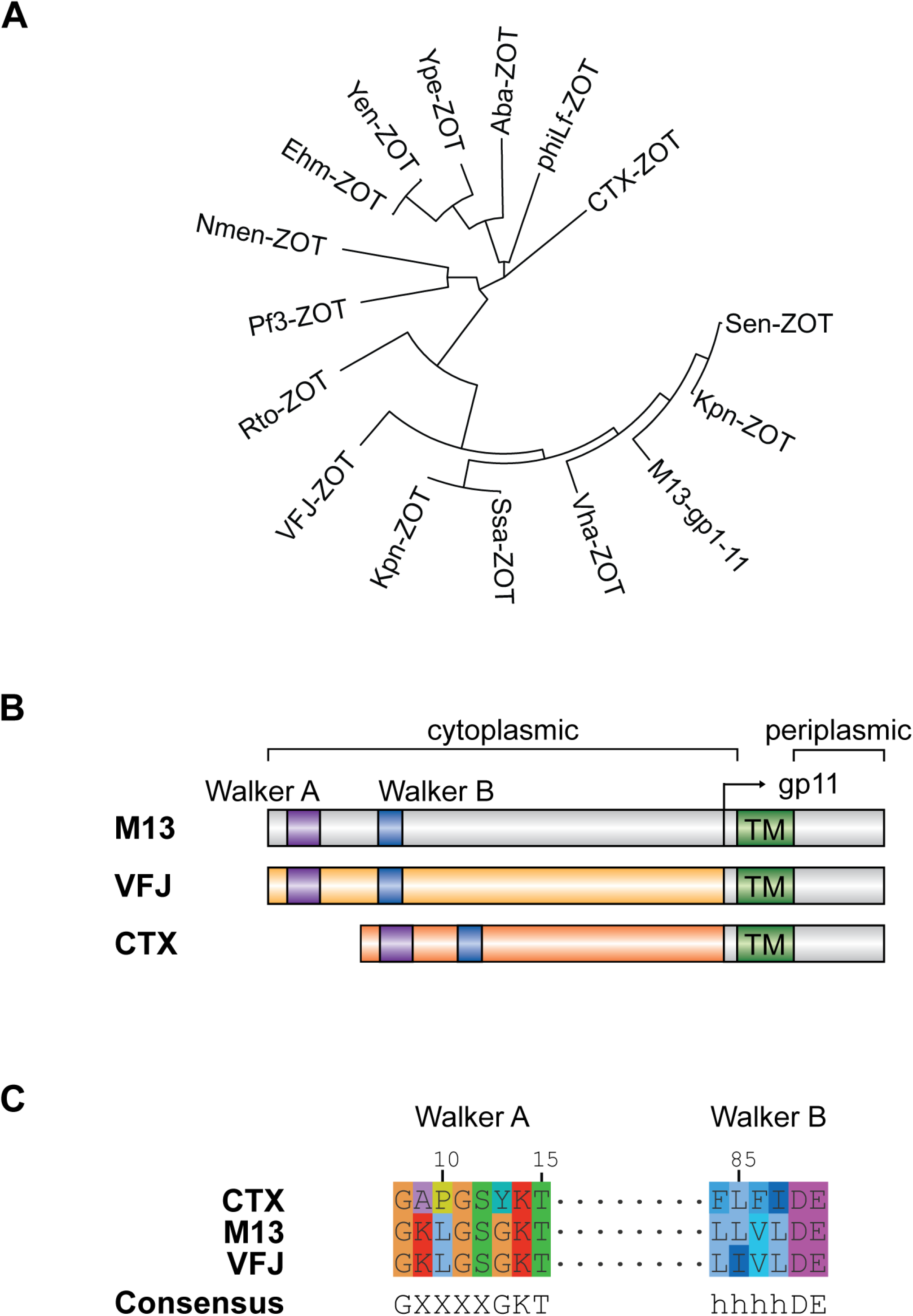
A) Circular phylogenetic tree comparing various Zot-like proteins from different organisms. Sen-: *Salmonella enterica*; Kpn-: *Klebsiella pneumoniae*; M13-: *Escherichia* virus M13; Vha-: *Virgibacillus halodenitrificans*; Ssa-: *Streptococcus sanguinis*; VFJ-: *Vibrio* virus VFJφ; Rto-: *Ruminococcus torques*; Pf3-: *Pseudomonas* virus Pf3; Nmen-: *Neisseria meningitidis*; Ehm-: *Enterobacter hormaechei*; Yen-: *Yersinia enterocolitica*; Ype-: *Yersinia pestis*; Aba-: *Acinetobacter baumannii*; phiLf-: *Xanthomonas* phage phiLf; CTX-: *Vibrio* virus CTXφ. B) Schematic representation of full length ZOT and ZOT-like proteins from M13, VFJϕ and CTXφ. ZOT-like g1p in both M13 and VFJϕ contain an internal ORF which encodes gene 11. In CTXφ, the start of gene 11 and its function are less clear. Purple box: Walker A motif. Blue box: Walker B motif. Arrow indicates internal open reading frame of g1p1. TM: transmembrane region. C) Amino acid alignment of Walker A and Walker B motifs in the ZOT protein of filamentous phages M13, VFJϕ and CTXφ. Numbers above the alignment indicate amino acid positions in M13 gene1. Consensus sequences are indicated below.

### Deviant Walker A and Walker B sequences in the CTXϕ Zot nucleotide binding pocket

The morphogenesis protein of the filamentous coliphage M13 consists of two parts, which are separated by a transmembrane domain. The cytoplasmic domain contains a Walker A and a Walker B motif, while the periplasmic domain is thought to form a continuous secretion tunnel together with the outer membrane protein g4p or p4, allowing the assembled phage filament to cross the cell envelope (Figure 1B). When comparing the putative molecular motor protein complex of filamentous phages from different species, we found that the Walker A and Walker B motifs do not necessarily follow the characteristic amino acid consensus sequence (Supplemental Figure 1). Walker A, also known as the P- or phosphate binding loop, frequently contains a leucine at position 10 (according to the g1p of the coliphage M13) in most g1p homologues analysed. This allows a putative helix to form that, together with the beta-sheet structure of the Walker B, allows the formation of a groove where ATP is able to bind prior to its hydrolysis. Several sequences however, display a proline at this position instead, which creates a kink in protein structures and is therefore helix-breaking. Somewhat surprising is the observation of a bulky tyrosine residue in position 13 in the case of the protein from the *Vibrio cholerae* phage CTXϕ. According to the classical Walker A motif, this residue is occupied by a conserved glycine, the smallest and most rotationally flexible residue, and no other amino acids have been reported so far (in all known ATPases, not only viral ones). *V. cholerae* is the host for several filamentous phages, not exclusively CTXϕ but also a phage called VFJϕ. The Walker A and B sequences of this phage are almost identical to the one from M13, with both, the P10 and the Y13 absent (Figure 1C). Walker B is described as four hydrophobic residues followed by a D and E. In all g1p sequences of the studied phages we found this to be the case. However, in the case of CTXϕ, two phenylalanine residues are present in position 84 and 86. Albeit hydrophobic, the 2 residues are considered rather bulky as the benzene group occupies a comparably large space. In all other cases, the correlating residues are leucine, isoleucine or valine residues.

### The introduction of deviant residues in Walker A and Walker B in M13 g1p abolished phage production

To understand the impact of the residues we observed in the putative nucleotide binding regions in CTXϕ g1p, we conducted a series of *in vivo* tests where we mutated one residue at a time. In our *in vivo* assay, phage production is assessed by making use of an amber mutant of phage M13 in which gene 1 is disrupted, unless it is present in a suppressor strain or complemented by a functional gene 1 *in trans*. We previously showed that such a complementation by a plasmid-encoded gene 1 is possible and allows the study of the impact of individual mutations in the gene ^22^. To investigate the impact of the homologue sequences observed in CTXϕ, we tested the complementation *in trans* for the individual and combined mutations in M13 g1p. The replacement of the catalytic lysine residue in position 14 abolished phage production and served as a control ^22^. In contrast, the mutation of a lysine residue in position 9 to an alanine had no impact on phage production (Figure 2A and B). However, replacing a leucine in position 10 with the proline found in CTXϕ g1p, reduced phage production to the same extent as the replacement of the catalytic lysine did, with low numbers of phages being produced due to reversion or translation errors. Similarly, mutating the glycine in position 13 to a tyrosine found in CTXϕ, was not tolerated and almost no phages were produced. Also, combining mutations or replacing the entire CTXϕ-like Walker A motif in M13 g1p were unable to rescue the function of the protein.

**Figure 2:**
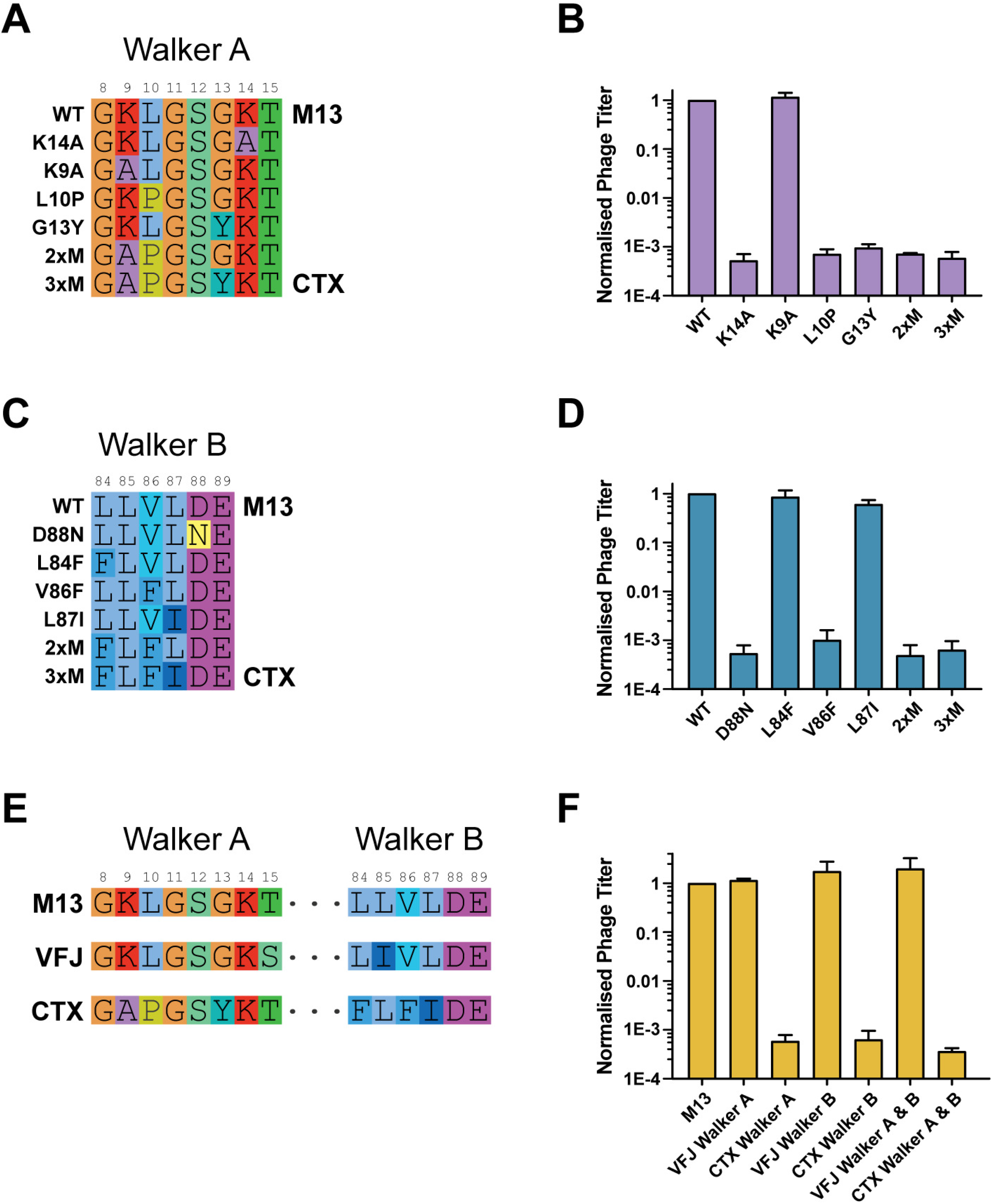
CTXφ Walker motifs result in dysfunctional protein. A & B) Residues in Walker A of M13 gene1 were mutated to CTXφ Walker A sequence and tested in *in vivo* complementation assays. C & D) Residues in Walker B of M13 gene1 were mutated to CTXφ Walker B sequence and tested in *in vivo* complementation assays. E & F) Residues in Walker A and B of M13 gene1 were mutated to VFJϕ and CTXφ Walker A and B sequence and tested in *in vivo* complementation assays. A,C,E) Amino acid alignment of Walker motifs. Numbers above the alignment indicate amino acid positions in M13 gene1. B,D,F) Phage titre from *in vivo* complementation assays with gene1 Walker motif mutants.

We then investigated the role of the mutations in Walker B by replacing the corresponding residues in M13 g1p with those found in CTXϕ. Here, we observed that the introduction of one of the two phenylalanine residues present in the *Vibrio* phage abolishes the function of the protein. While the L84F mutation has no impact on the function of the protein, assessed by the production of phages, the mutation V86F, resulted in a loss of function of g1p (Figure 2C and D). Again, combined mutations as well as the replacement of the entire Walker B -from M13 to CTXϕ - did not restore the function of the protein.

As Walker A and Walker B form a binding pocket for ATP in ATPases, we also investigated the mutations found in CTXϕ by introducing them into both motifs in M13 g1p and determined if they were able to restore protein function. Here we observed a severe reduction in the number of phages, indicating that the mutated protein is rendered non-functional by the amino acid substitutions (Figure 2E and F). As *V. cholerae* is the host of several filamentous phages, we questioned whether the gene1 homologue from *Vibrio cholerae* phage VFJϕ has a functional ATPase. To this end, we mutated both motifs in M13 g1p to the sequence found in VFJϕ and found that phage titres were unaffected regardless of the mutations made in individual motifs or combined, indicating that bacteriophage VFJϕ has a functional ATPase in the gene1 homologue (Figure 2E and F).

### VFJϕ Zot is able to mediate CTXϕ phage assembly in Vibrio cholerae

Since *V. cholerae* is the host of several filamentous phages and we found that VFJϕ-Zot encodes a functional ATPase *in vivo*, we asked whether it was possible for VFJϕ-Zot to mediate the assembly of CTXϕ. In order to address this question, we first established the assay to study VFJϕ and CTXϕ *in vivo* as the common method of determining plaque forming units by counting plaques is not a reliable approach here; phages VFJϕ and CTXϕ were previously shown to produce very few plaques and hence the chosen method of detection was qPCR ^26^. In order to validate the qPCR approach for detecting phages, we made use of M13. *Gene 3* coding for the receptor-binding protein g3p was used for amplification and was found to correlate with the number of phages determined by counting PFUs on bacterial culture plates (Supplementary Figure 2). As we also wanted to quantify the number of phages produced per cell, we used a set of primers to detect the ribosomal 16S gene in *V. cholerae*, following a previously established protocol ^26^.

We then conducted *in vivo* complementation assays using a *Vibrio cholerae* strain that contains the CTXϕ (but not the VFJϕ) prophage genome but has the cholera toxin and the Zot genes deleted by genetic engineering (ΔZot). The Zot gene from either VFJϕ or CTXϕ were then cloned under the control of an arabinose inducible promotor. As a negative control, the plasmid backbone that does not contain any gene was used. The constructs were introduced into the *V. cholera* ΔZot strain and the number of phages produced was quantified by qPCR. Using this method, the number of amplicons in the presence of CTXϕ-Zot remained low as in the case of the control plasmid. However, when VFJϕ-Zot was introduced into the *V. cholera* ΔZOT strain, we were able to detect phages (Figure 3). Phage production in our assay was comparably low possibly due to low prophage induction efficiency despite the presence of the inducer mitomycin C. Regardless of the low number, our data supports the hypothesis that CTXϕ-Zot is an inefficient or inactive assembly protein and the CTXϕ phage is able to “hijack” the protein from phage VFJϕ for its assembly.

**Figure 3:**
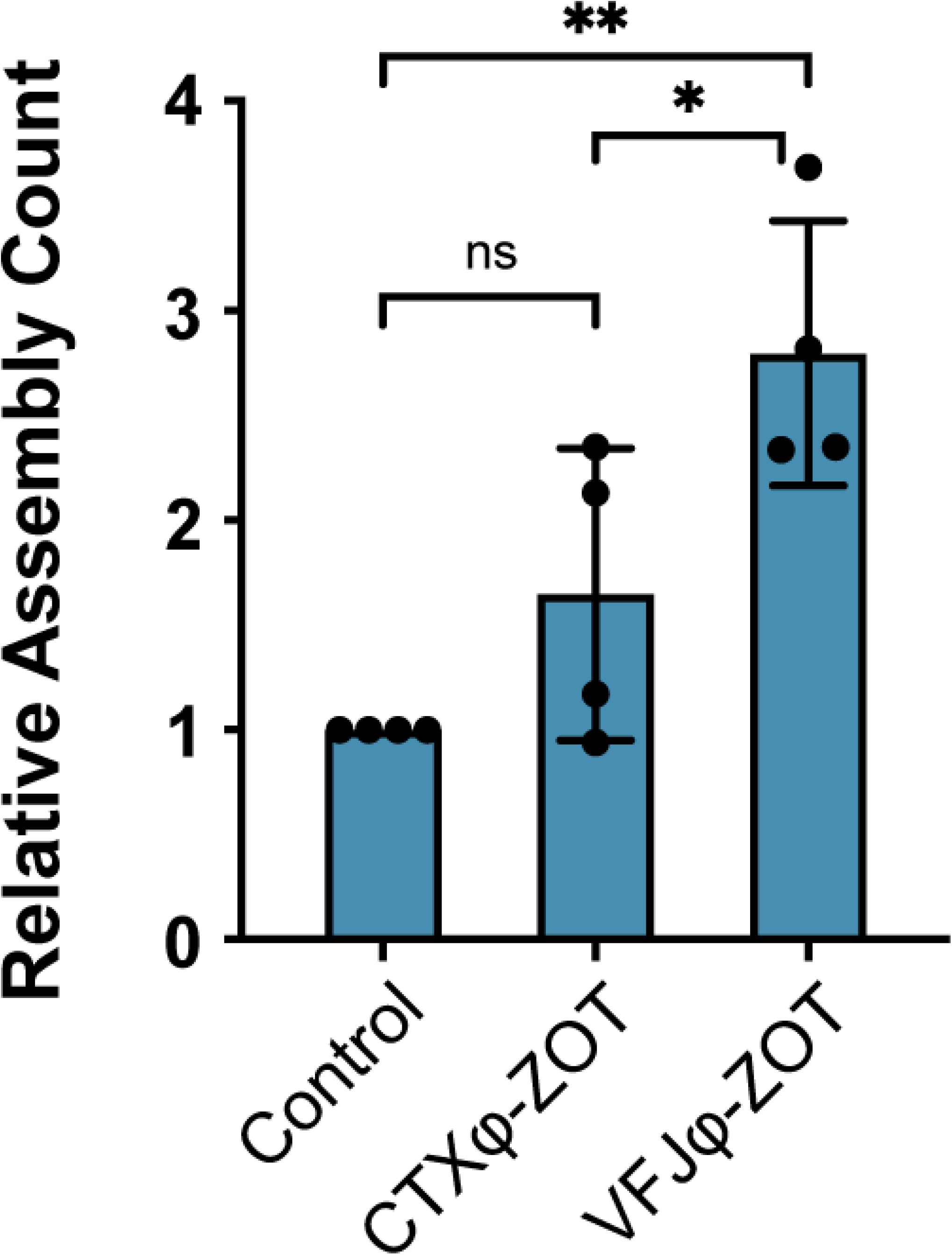
CTXφ-ZOT is dysfunctional *in vivo*. Expression of CTXφ-ZOT in *V. cholera* ΔZOT strain results in as few phages produced as seen in control. Expression of VFJφ-ZOT facilitates phage production. Control: Parent plasmid without ZOT gene.

### A highly conserved glutamine in g1p homologs is not conserved in CTXϕ but is essential for assembly

When aligning sequences of g1p homologs from several organisms, including *Vibrio cholerae, Escherichia coli, Acinetobacter baumannii, Neisseria meningitidis, Klebsiella pneumoniae* and others, we found a single highly conserved residue in addition to the Walker motifs. Among all investigated filamentous phage assembly proteins we found Q126 to be conserved, with the exception of the protein in CTXϕ; here, a proline can be found in the corresponding position (Figure 4A). To test if Q126 can be replaced by a proline or other residues, we created single point mutations and tested them in the M13 *in vivo* complementation assay. Our results show that in M13, g1p Q126P is not functional as is the replacement of glutamine by asparagine, glutamic acid and lysine among many others (Figure 4B). Surprisingly, the only mutation we found to be tolerated, was a replacement with methionine, which shares the same length but not the same charge with glutamine. At present, the function of this residue remains speculative.

**Figure 4:**
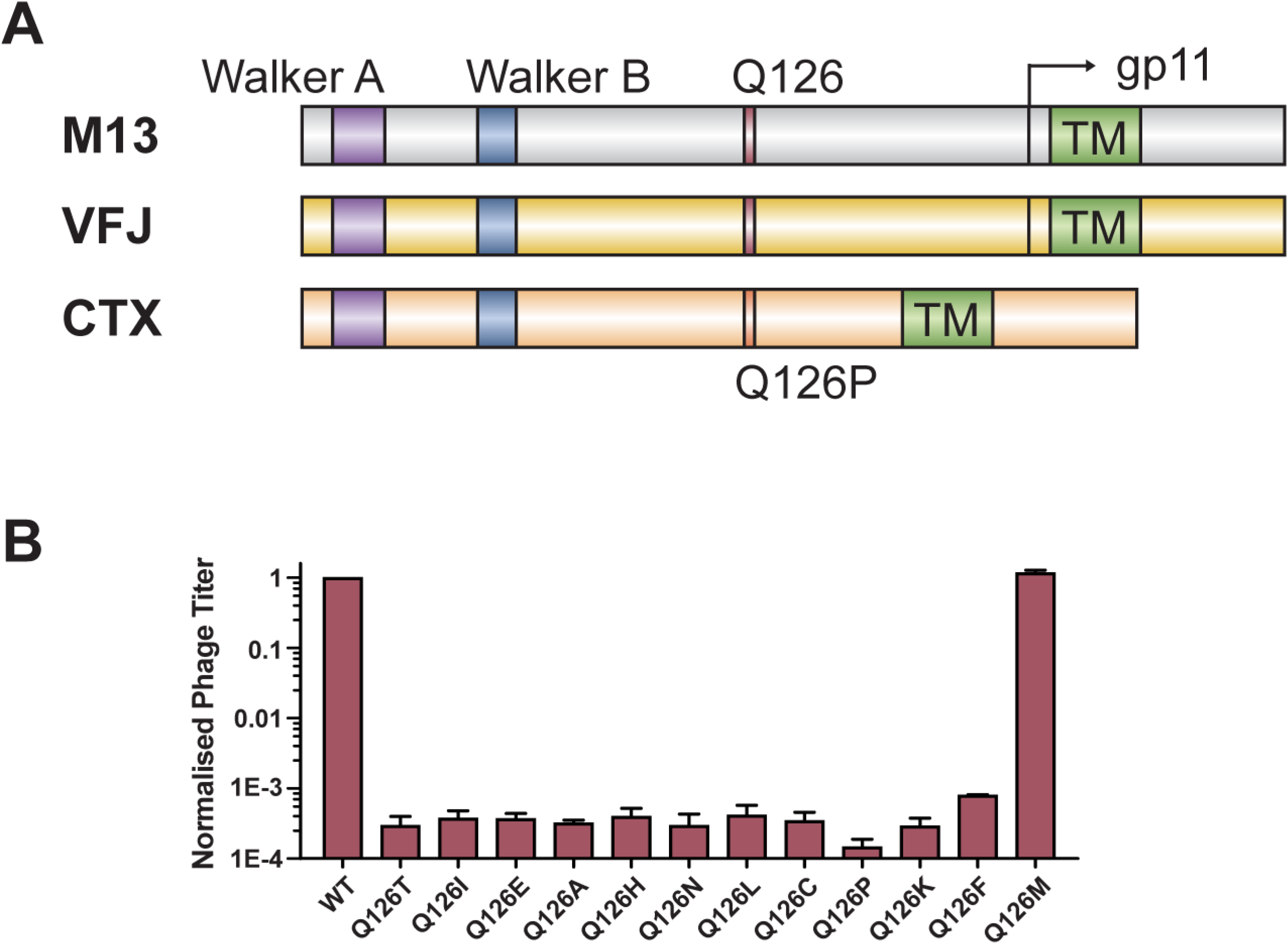
Residue Q126 is important for protein function. A) Schematic representation of the M13 gene1 and homologues in VFJϕ and CTXφ, with position of the Q residue included. B) Phage titre from *in vivo* complementation assays with gene1 Q126 mutants.

### Structure prediction and molecular dynamics simulation indicate a regulatory role of Q126 for ATP binding

As the role of the glutamine (Q126) in the hydrolysis of ATP by functional assembly proteins is unclear, we resorted to structural biology using AlphaFold2 to predict the structures of M13 g1p, VFJϕ-Zot and CTXϕ-Zot. The predicted structures show that M13 g1p, VFJϕ-Zot and CTXϕ-Zot share a similar topology (Figure 5A). The cytoplasmic domains among the 3 proteins are highly similar however, CTXϕ-Zot is slightly more distinct from M13 g1p and VFJϕ-Zot, as indicated by the root mean squared deviations (RMSD) of VFJϕ-Zot and CTXϕ-Zot from M13 g1p (0.1 nm and 0.38 nm, respectively).

**Figure 5:**
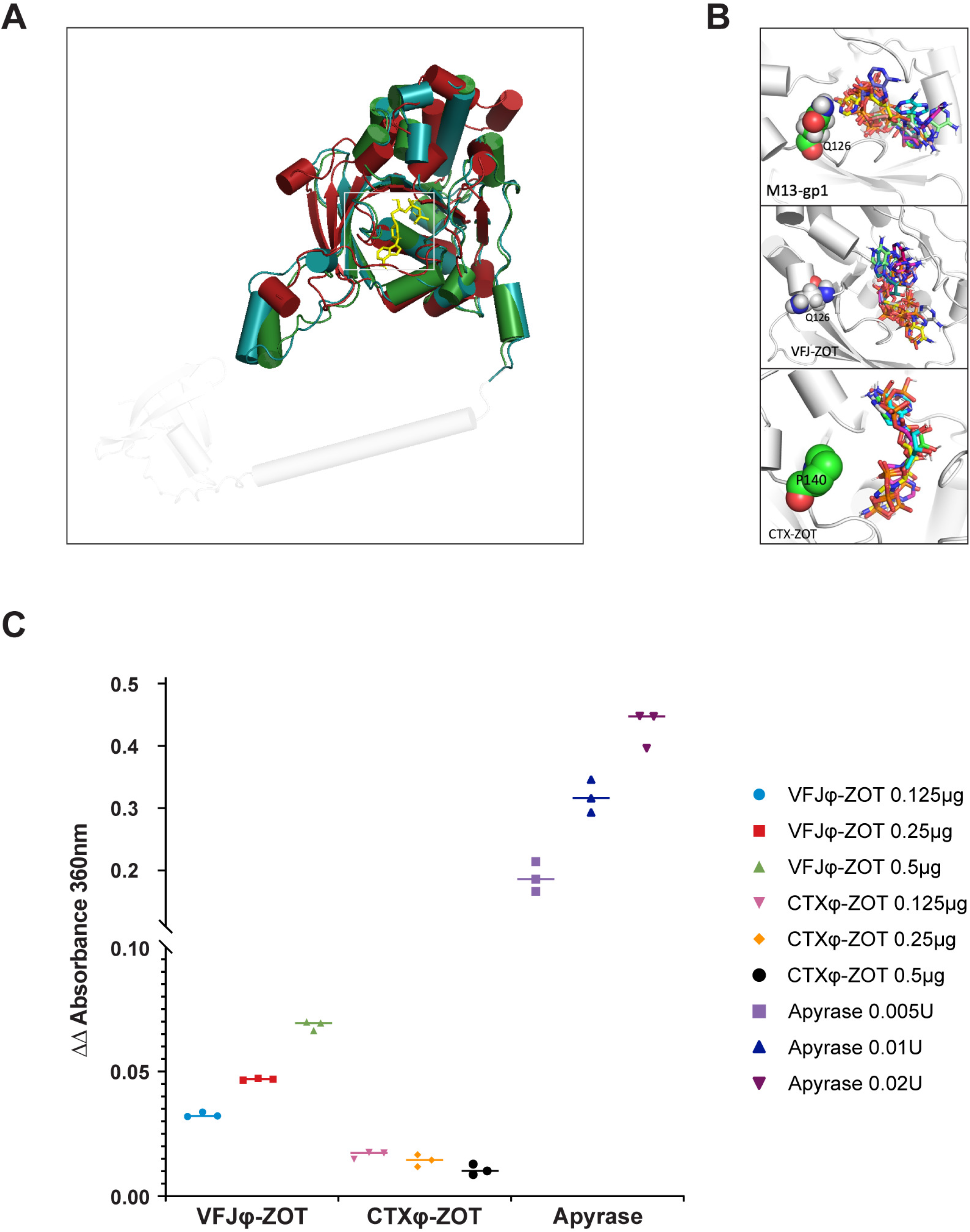
Structures of M13-g1p, VFJϕ-Zot and CTXϕ-Zot predicted by AlphaFold2. A) Five models were predicted for each protein. The models are represented as cartoons coloured in blue to red from the N-terminal region to the C-terminal region. B) Molecular docking of ATP within M13 g1p, VFJϕ Zot and CTXϕ Zot. The top six structures are represented. Note that different from M13 g1p and VFJϕ Zot, the residue at the corresponding position of Q126 is P140 in CTXϕ Zot.

Interestingly, in the predicted structure of M13 g1p, Q126 is positioned in close proximity to the ATP-binding pocket (Figure 5B). We speculated that Q126 may play an active role in ATP binding and release in regulating phosphate hydrolysis. To test this hypothesis, we performed molecular docking on the groove formed by residues from the Walker motifs and also all-atom molecular dynamics (MD) simulation of M13 g1p based on the docked structures. In the MD simulation, we indeed observed that Q126 could form extensive hydrogen bonding interactions with the nucleotide (Figure 5B and Movie in SI), thus supporting a crucial role in regulating ATP hydrolysis. This may also explain why mutations are not well tolerated in the protein including the proline residue found in the corresponding position of Q126 in CTXϕ-Zot. Interestingly, in the structures, we observed a similar groove in all three proteins allowing the binding of ATP molecules, assessed by molecular docking. However, the binding modes of ATP in CTXϕ Zot are distinctly different from M13 g1p and VFJϕ Zot. Aside from Q126, we also observed other residues, such as S12, G13, K14, T15 in Walker A motif and R148, H175, T206, K207 in the loop region, to form stable interactions with ATP, suggesting that they may contribute to ATP binding (Supplementary Figure 3).

### The cytoplasmic domain of VFJϕ-Zot is able to hydrolyse ATP in vitro, while CTXϕ-Zot shows no activity

To test if the CTXϕ-Zot or VFJϕ-Zot proteins show ATPase activity *in vitro*, we cloned and expressed the cytoplasmic domains as MBP-fusion proteins in *E. coli* and purified them to homogeneity. The proteins where then tested for their ability to hydrolyse the nucleotide *in vitro* in an absorbance-based assay. While the Zot-like protein from VFJϕ clearly showed ATPase activity in a concentration dependent manner, we were unable to detect any ATP hydrolysis in the case of the CTXϕ-Zot protein (Figure 5C and Supplementary Figure 4). However, often additional molecules such as substrate or co-factors are required for proteins to show (full) activity if they are tightly controlled. With this caveat in mind, the lack of activity of CTXϕ-Zot in our assay indicates that the protein might be an inefficient ATPase, or one that has lost its ability to hydrolyse ATP entirely.

### Divergence of Walker motifs in Vibrio genomes indicate that the CTXϕ Zot-like Walker sequences are prevalent in Vibrio cholerae but not in other Vibrio species

Using previously developed HMM profiles ^37^, we identified 1,203 g1p-like proteins across 1,122 genomes of *Vibrio cholerae*. After aligning these proteins, we could categorise the proteins into three main “variants”, i.e. combinations of mutations: (1) The CTX-like, containing three non-functional mutations in the Walker motifs and a Q126P mutation, (2) the functional M13/VFJ-like including a Q126 residue, and (3) a “hybrid” category which contains one deleterious mutation in the Walker A or B motif and the Q126P mutation. One more category called “atypical” includes all other proteins that were detected by the algorithm with partially large variations in sequence. Here, in some instances the Walker A follows the GxxxxGKT/S consensus, yet with a proline in position 10 (47% of the “atypical” sequences). Others do not conform with the consensus motif albeit the catalytic lysine residue is present. Most of the proteins in *Vibrio cholerae* are CTXϕ-like (842) or hybrid (263), and only 9 were clearly M13/VFJϕ-like, while the remaining 89 were g1p candidates with atypical mutations (Supplemental table 3).

When analyzing putative filamentous prophages in *Vibrio sp*. excluding *V. cholerae*, we found that 1,607, of 5,117 (~26%) contained M13/VFJϕ-like sequences, while only 7 displayed a combination of mutations strictly identical to CTX (0.001%). These were all found in the *Vibrio mimicus* species, which is known to sometimes encode cholera-like toxins ^38^. However, a number of pI-like proteins found in other vibrio species than *V. cholera* included some, but not all, of the CTX-like deleterious mutations. For instance, we observed 187 instances of L10P, 5,143 instances of G13Y, 112 instances of V86F, and 3,731 instances of Q126P. This suggests that pI-like ATPases in *Vibrio* can accumulate deleterious mutations relatively easily, most likely because of the possibility of complementation by an intact pI-like ATPase from a co-infecting filamentous phage.

## Discussion

*Vibrio cholerae* is the host for many phages and several of them are filamentous ^16,39–42^. In this work we studied the Zonula occludens toxin (Zot) of the *Vibrio cholerae* phages VFJϕ and CTXϕ and compared it to the highly homologous filamentous phage assembly protein found in M13. CTXϕ is well known for its potential for toxigenic conversion, i.e. the conversion of *Vibrio cholerae* strains into virulent ones that express the phage-encoded cholera toxin, making them more pathogenic. This phage has been shown to encode additional toxins, one of which is Zot. Zot has been reported to create lesions in tissue, specifically in the *zonlula occludens*, when investigating *V. cholerae* culture supernatants or the recombinant protein Zot ^25,43,44^.

Zot contains regions that align with high homology to the assembly (or morphogenesis) protein g1p of the coliphage M13, also known as p1, pI, g1p. Our previous work on the gene1 encoded assembly complex of the coliphage M13 demonstrated that both Walker motifs are essential for phage assembly ^22^. Making use of this system, we tested the deviant motifs found in CTXϕ in our *in vivo* assay.

While Walker A and Walker B motifs remain identifiable when aligning related sequences to each other, Walker A variants have been identified by aligning the large terminase proteins of lytic bacteriophages ^45^. However, the consensus sequence for Walker A can be defined as S/GxxxxGKT/S (with x being any residue). In our study, a deviation from the classical Walker A motif seems apparent: A small and flexible glycine residue in position 13 (numbering according to g1p of M13) preceding the catalytic lysine, present in all Walker A motifs reported thus far, is replaced by the large, aromatic side chain of a tyrosine in CTXϕ Zot. In addition, a helix-breaking proline is found in position 10 where a leucine is found in most g1p homologs. In Walker B of CTXϕ Zot, two phenylalanine residues are found where more commonly less bulky hydrophobic residues are present, which precede the characteristic DE dyad found in the motif. Our extensive set of experiments including single point mutations, their combination and also domain swapping experiments of the protein from CTXϕ (data not shown), indicate that CTXϕ Zot is not a functional ATPase. Hence, we propose that in order to produce CTXϕ phage particles, a co-infection with a second filamentous phage -such as VFJϕ-is required. This might allow CTXϕ to exploit the assembly complex from another phage, a phenomenon reported for other phages ^14^. A major discrepancy between our finding and a previous report is the publication by Faruque, S.M. et al. (2002), describing the biogenesis of filamentous phage particles found in *Vibrio cholerae*. The authors stipulate that the genetic element RS1 which can be encapsidated into a filamentous phage particle, requires another phage ^17^. RS1 itself does not code for any morphogenesis or coat proteins and thus would rely on a phage with functional genes, including g1p. While the experimental evidence is extensive and the conclusion of the publication sound, no whole genome sequence of the tested bacterial strains is available, which allows the possibility that a second filamentous phage genome may be present in the tested isolates.

To explore the question of whether a “borrowed” assembly protein could facilitate producing phage particles, we made use of a strain that contains only one filamentous phage, CTXϕ, in which the Zot protein has been genetically removed. We determined the number of phage particles produced in the strain when complemented with the Zot gene *in trans*, or the Zot-like gene from VFGϕ which is highly similar to the assembly gene from M13. While induction and viral production conditions in this plasmid complementation experiments might not be optimal, we could establish that no phage particles were produced when the CTXϕ ZOT gene was introduced on a plasmid into the host, similar to the “empty” control plasmid. In contrast, a plasmid encoding the VFJϕ ZOT-like gene resulted in the production of phage particles albeit at a low number, indicating that the expressed protein has the ability to encapsidate the DNA of CTXϕ by assembling the coat proteins of the virus around the genome.

The ATPase activity we show here is the first demonstration of such activity in an *in vitro* assay using the purified Zot protein. A previous attempt conducted by Schmidt et al. ^44^ obtained purified protein by denaturing the polypeptide using a chaotropic agent (guanidine HCl), which leads to protein unfolding and the spontaneous refolding in physiological buffers is not guaranteed. Especially for kinases and ATPases, refolding from denatured proteins can be challenging as often molecular chaperones are required for folding ^27^. To address the question of whether the proteins from the two *Vibrio* phages have the ability to hydrolyse ATP, we purified the cytoplasmic domains of the proteins from both phages as MBP-fusion proteins, as the full length proteins could not be captured on the column or showed toxicity towards *E. coli* during expression. Consistent with the previously reported study by Schmidt et al, the ATPase domain of CTXϕ Zot shows no ATPase activity. However, the corresponding domain from the VFJϕ phage clearly has the ability to hydrolyse the nucleotide. The observed hydrolysis rate of the protein is not extensive compared to the control, the enzyme apyrase, possibly due to being tightly controlled by co-factors such as host- and viral proteins, and the phage DNA. The *in vitro* assay using the purified proteins provides additional evidence to hypothesize that CTXϕ Zot is a protein that has gained another, possibly more important, function during evolution. As the phage might be able to be propagated by using another phage’s assembly complex (as established for other filamentous phages), the CTXϕ Zot could develop to become a secondary toxin in addition to the cholera toxin CTX. Additional evidence that CTXϕ Zot may have evolved “away” from a motor protein to become a toxin is the sequence that the protein displays in the periplasmic region in case of M13 g1p and other filamentous phages. Here, the sequence appears to have almost no homology and little structural similarity (Supplementary Figure 1).

If CTXϕ has a dysfunctional motor protein (yet a functional accessory toxin), how does the virus replicate? According to a previous study *Vibrio* strains often contain more than one filamentous phage type, which would allow them to exploit the assembly machine of other phages. As the report was published in 2011, we employed a recently established algorithm by Simon Roux et al ^37^, to analyse how many filamentous prophages are present per genome in *V. cholerae* strains. We found that ~68% (n = 1,040) contain one filamentous prophage and 4% (n = 66) have two or more prophages, while none could be detected in 423 of 1,529 (~28%) strains (Supplementary Figure 4B). While this is significantly more than the 45% reported in a recent study which however analysed the sequences of all *Vibrio* species for Inoviridae prophage sequences ^41^, our findings also indicate that CTXϕ would in most cases not be transmitted horizontally, should our interpretation of the findings in this study be accurate.

Our analysis of *Vibrio cholera* genomes might suggest that the CTXϕ-like protein has been “domesticated” for use as a toxin and is probably not able to function as a pI-like protein for inovirus assembly. Proteins encoding a “hybrid” sequence contain a tyrosine in Walker A where a glycine should be present, which in our *in vivo* test model would result in a non-functional protein. At present, it is unclear if such proteins are possibly coding for functional ATPases, which would be a novelty in the field of understanding ATP-binding and hydrolysis mechanisms. However, during evolution prophages can undergo genetic alterations rendering them non-functional and unable to generate functional phage particles, while genes that provide an evolutionary advantage to the host such as toxins are more likely to be conserved.

Indication that the proteins with “deviant motifs” might be able to bind ATP and possibly hydrolyse the nucleotide is given by the AlphaFold2 structure prediction, molecular docking and all-atom MD simulation. Although all three proteins, CTXϕ-Zot, VFJϕ-Zot and M13-g1p, show a conserved ATP binding pocket in a structurally similar cytoplasmic domain, the residue Q126 in M13-g1p, observed to form extensive interactions with the nucleotide in the MD simulation (Supplementary movie MD3) is replaced by a proline residue in the corresponding position in CTXϕ-Zot. This mutation in CTXϕ-Zot may reduce the binding affinity with ATP by breaking the interactions, and therefore abolish ATP hydrolysis, despite the overall structure still being conserved.

Based on the data we obtained from our *in vivo* tests, bioinformatic analyses and *in vitro* enzymatic assays, we argue that the ZOT gene in *V. cholerae* phage CTXϕ has evolutionarily lost its original function, i.e. to facilitate the assembly of the phage particle in the membrane, and now fulfils the role of an auxiliary toxin during the infection process for its host.

## Acknowledgments

We would like to thank Prof. Menghua Yang (ZAFU, Hangzhou, China) for providing us with El Tor *V. cholerae* strain C6706, and for her advice on working with the pathogen, for which we would like to thank Prof. Julia Fritz-Steuber (University of Hohenheim, Germany) as well. This work was supported by a grant from the National Natural Science Foundation of China (32011530116). And the simulations were supported by Information Technology Center and State Key Lab of CAD&CG, Zhejiang University. The work conducted by the U.S. Department of Energy Joint Genome Institute, a DOE Office of Science User Facility, is supported under Contract No. DE-AC02-05CH11231.

## Author Contributions

B.L. and S.L. conceived the study, B.L. and M.L. designed the experiments; M.L., B.L., X.H., S.R. and S.L. performed the experiments or analysed the data; B.L prepared the figures; S.L. and B.L. wrote the paper. All authors agreed to the final version of the manuscript.

## References

1 Boyd, E. F. & Brussow, H. Common themes among bacteriophage-encoded virulence factors and diversity among the bacteriophages involved. Trends Microbiol 10, 521–529, doi:10.1016/s0966-842x(02)02459-9 (2002).

2 Herold, S., Karch, H. & Schmidt, H. Shiga toxin-encoding bacteriophages--genomes in motion. Int J Med Microbiol 294, 115–121, doi:10.1016/j.ijmm.2004.06.023 (2004).

3 Tinsley, C. R., Bille, E. & Nassif, X. Bacteriophages and pathogenicity: more than just providing a toxin? Microbes Infect 8, 1365–1371, doi:10.1016/j.micinf.2005.12.013 (2006).

4 Jamet, A. et al. A widespread family of polymorphic toxins encoded by temperate phages. BMC Biol 15, 75, doi:10.1186/s12915-017-0415-1 (2017).

5 Loh, B., Kuhn, A. & Leptihn, S. The fascinating biology behind phage display: filamentous phage assembly. Mol Microbiol 111, 1132–1138, doi:10.1111/mmi.14187 (2019).

6 Sweere, J. M. et al. Bacteriophage trigger antiviral immunity and prevent clearance of bacterial infection. Science 363, doi:10.1126/science.aat9691 (2019).

7 Wendling, C. C., Refardt, D. & Hall, A. R. Fitness benefits to bacteria of carrying prophages and prophage-encoded antibiotic-resistance genes peak in different environments. Evolution 75, 515–528, doi:10.1111/evo.14153 (2021).

8 Waldor, M. K. & Mekalanos, J. J. Lysogenic conversion by a filamentous phage encoding cholera toxin. Science 272, 1910–1914, doi:10.1126/science.272.5270.1910 (1996).

9 Kuhn, A. & Leptihn, S. Helical and filamentous phages. (2019).

10 Roux, S. et al. Author Correction: Cryptic inoviruses revealed as pervasive in bacteria and archaea across Earth’s biomes. Nat Microbiol 5, 527, doi:10.1038/s41564-020-0681-5 (2020).

11 Rice, S. A. et al. The biofilm life cycle and virulence of Pseudomonas aeruginosa are dependent on a filamentous prophage. ISME J 3, 271–282, doi:10.1038/ismej.2008.109 (2009).

12 Burgener, E. B. et al. Filamentous bacteriophages are associated with chronic Pseudomonas lung infections and antibiotic resistance in cystic fibrosis. Sci Transl Med 11, doi:10.1126/scitranslmed.aau9748 (2019).

13 Shapiro, J. W. & Putonti, C. UPPhi phages, a new group of filamentous phages found in several members of Enterobacteriales. Virus Evol 6, veaa030, doi:10.1093/ve/veaa030 (2020).

14 Rakonjac, J., Bennett, N. J., Spagnuolo, J., Gagic, D. & Russel, M. Filamentous bacteriophage: biology, phage display and nanotechnology applications. Current issues in molecular biology 13, 51–76 (2011).

15 Rubin, E. J., Lin, W., Mekalanos, J. J. & Waldor, M. K. Replication and integration of a Vibrio cholerae cryptic plasmid linked to the CTX prophage. Mol Microbiol 28, 1247–1254, doi:10.1046/j.1365-2958.1998.00889.x (1998).

16 Davis, B. M., Moyer, K. E., Boyd, E. F. & Waldor, M. K. CTX prophages in classical biotype Vibrio cholerae: functional phage genes but dysfunctional phage genomes. J Bacteriol 182, 6992–6998, doi:10.1128/JB.182.24.6992-6998.2000 (2000).

17 Faruque, S. M. et al. RS1 element of Vibrio cholerae can propagate horizontally as a filamentous phage exploiting the morphogenesis genes of CTXphi. Infect Immun 70, 163–170, doi:10.1128/IAI.70.1.163-170.2002 (2002).

18 Faruque, S. M. et al. CTXphi-independent production of the RS1 satellite phage by Vibrio cholerae. Proc Natl Acad Sci U S A 100, 1280–1285, doi:10.1073/pnas.0237385100 (2003).

19 Hassan, F., Kamruzzaman, M., Mekalanos, J. J. & Faruque, S. M. Satellite phage TLCphi enables toxigenic conversion by CTX phage through dif site alteration. Nature 467, 982–985, doi:10.1038/nature09469 (2010).

20 Krupovic, M., Prangishvili, D., Hendrix, R. W. & Bamford, D. H. Genomics of bacterial and archaeal viruses: dynamics within the prokaryotic virosphere. Microbiol Mol Biol Rev 75, 610–635, doi:10.1128/MMBR.00011-11 (2011).

21 Rapoza, M. P. & Webster, R. E. The products of gene I and the overlapping in-frame gene XI are required for filamentous phage assembly. J Mol Biol 248, 627–638, doi:10.1006/jmbi.1995.0247 (1995).

22 Loh, B., Haase, M., Mueller, L., Kuhn, A. & Leptihn, S. The Transmembrane Morphogenesis Protein gp1 of Filamentous Phages Contains Walker A and Walker B Motifs Essential for Phage Assembly. Viruses 9, doi:10.3390/v9040073 (2017).

23 Koonin, E. V., Tatusov, R. L. & Rudd, K. E. Sequence similarity analysis of Escherichia coli proteins: functional and evolutionary implications. Proc Natl Acad Sci U S A 92, 11921–11925, doi:10.1073/pnas.92.25.11921 (1995).

24 Uzzau, S., Cappuccinelli, P. & Fasano, A. Expression of Vibrio cholerae zonula occludens toxin and analysis of its subcellular localization. Microb Pathog 27, 377–385, doi:10.1006/mpat.1999.0312 (1999).

25 Di Pierro, M. et al. Zonula occludens toxin structure-function analysis. Identification of the fragment biologically active on tight junctions and of the zonulin receptor binding domain. J Biol Chem 276, 19160–19165, doi:10.1074/jbc.M009674200 (2001).

26 Takahashi, N. et al. Lethality of MalE-LacZ hybrid protein shares mechanistic attributes with oxidative component of antibiotic lethality. Proc Natl Acad Sci U S A 114, 9164–9169, doi:10.1073/pnas.1707466114 (2017).

27 Sievers, F. et al. Fast, scalable generation of high-quality protein multiple sequence alignments using Clustal Omega. Mol Syst Biol 7, 539, doi:10.1038/msb.2011.75 (2011).

28 Letunic, I. & Bork, P. Interactive Tree Of Life (iTOL) v5: an online tool for phylogenetic tree display and annotation. Nucleic Acids Res 49, W293–W296, doi:10.1093/nar/gkab301 (2021).

29 Trott, O. & Olson, A. J. AutoDock Vina: improving the speed and accuracy of docking with a new scoring function, efficient optimization, and multithreading. J Comput Chem 31, 455–461, doi:10.1002/jcc.21334 (2010).

30 Jo, S., Kim, T., Iyer, V. G. & Im, W. CHARMM-GUI: a web-based graphical user interface for CHARMM. J Comput Chem 29, 1859–1865, doi:10.1002/jcc.20945 (2008).

31 Lomize, A. L., Pogozheva, I. D. & Mosberg, H. I. Anisotropic solvent model of the lipid bilayer. 1. Parameterization of long-range electrostatics and first solvation shell effects. J Chem Inf Model 51, 918–929, doi:10.1021/ci2000192 (2011).

32 Huang, J. et al. CHARMM36m: an improved force field for folded and intrinsically disordered proteins. Nat Methods 14, 71–73, doi:10.1038/nmeth.4067 (2017).

33 Abraham, M. J. et al. GROMACS: High performance molecular simulations through multi-level parallelism from laptops to supercomputers. SoftwareX 1, 19–25 (2015).

34 DeLano, W. L. Pymol: An open-source molecular graphics tool. CCP4 Newsl. Protein Crystallogr 40, 82–92 (2002).

35 Kitts, P. A. et al. Assembly: a resource for assembled genomes at NCBI. Nucleic Acids Res 44, D73–80, doi:10.1093/nar/gkv1226 (2016).

36 Eddy, S. R. Accelerated Profile HMM Searches. PLoS Comput Biol 7, e1002195, doi:10.1371/journal.pcbi.1002195 (2011).

37 Roux, S. et al. Cryptic inoviruses revealed as pervasive in bacteria and archaea across Earth’s biomes. Nat Microbiol 4, 1895–1906, doi:10.1038/s41564-019-0510-x (2019).

38 Spira, W. M. & Fedorka-Cray, P. J. Purification of enterotoxins from Vibrio mimicus that appear to be identical to cholera toxin. Infect Immun 45, 679–684, doi:10.1128/iai.45.3.679-684.1984 (1984).

39 Campos, J. et al. VGJ phi, a novel filamentous phage of Vibrio cholerae, integrates into the same chromosomal site as CTX phi. J Bacteriol 185, 5685–5696, doi:10.1128/JB.185.19.5685-5696.2003 (2003).

40 Canchaya, C., Proux, C., Fournous, G., Bruttin, A. & Brussow, H. Prophage genomics. Microbiol Mol Biol Rev 67, 238–276, table of contents, doi:10.1128/MMBR.67.2.238-276.2003 (2003).

41 Castillo, D. et al. Widespread distribution of prophage-encoded virulence factors in marine Vibrio communities. Sci Rep 8, 9973, doi:10.1038/s41598-018-28326-9 (2018).

42 Hay, I. D. & Lithgow, T. Filamentous phages: masters of a microbial sharing economy. EMBO Rep 20, doi:10.15252/embr.201847427 (2019).

43 Fasano, A. et al. Vibrio cholerae produces a second enterotoxin, which affects intestinal tight junctions. Proc Natl Acad Sci U S A 88, 5242–5246, doi:10.1073/pnas.88.12.5242 (1991).

44 Schmidt, E., Kelly, S. M. & van der Walle, C. F. Tight junction modulation and biochemical characterisation of the zonula occludens toxin C-and N-termini. FEBS Lett 581, 2974–2980, doi:10.1016/j.febslet.2007.05.051 (2007).

45 Mitchell, M. S. & Rao, V. B. Novel and deviant Walker A ATP-binding motifs in bacteriophage large terminase-DNA packaging proteins. Virology 321, 217–221, doi:10.1016/j.virol.2003.11.006 (2004).

